# Redox Signaling Mediates Differentiation of Adipose Progenitors in Response to Inflammatory Cytokines in the Adipose Tissue Secretome

**DOI:** 10.1101/2024.09.27.615547

**Authors:** Jessica L. Wager, Larissa G. Baker, Taylor B. Scheidl, Sophie Z. Yonan, Pina Colarusso, Antoine Dufour, Jennifer A. Thompson

**Author notes:** <u>Corresponding Author</u>: Jennifer Thompson PhD, Associate Professor, Cumming School of Medicine, University of Calgary, HMRB 78 3330 Hospital Dr. NW, Calgary, Alberta, Canada T2N 4N1.

## Abstract

Adipogenesis, the terminal differentiation of adipose progenitor cells (APCs), is critical in maintaining the functional integrity of adipose depots under obesogenic conditions. It is thought that a signal arising from the adipose microenvironment triggers APC differentiation; yet the identity and source of the signal remains unknown. This study sought to uncover the signal responsible for activating adipogenesis. Redox signaling was shown to influence adipogenesis in primary murine APCs treated with pharmacologic agents to manipulate the levels of reactive oxygen species (ROS). Increased generation of superoxide (O_2_^-^) and hydrogen peroxide (H_2_O_2_) *via* redox cyclers amplified both early and late APC differentiation, while ROS scavengers and antioxidants blunted differentiation. The impact of specifically targeting H_2_O_2_ with the antioxidant, catalase, or a catalase inhibitor, was restricted to lipid accumulation in late adipogenesis. Protein was concentrated from conditioned media of adipose tissue explants cultured *ex vivo* to capture signals within the adipose secretome. Differentiation was enhanced in APCs cultured in the presence of the adipose secretome, an effect that was diminished with scavenging of ROS and amplified when the secretome was collected from mice fed a high fat diet for 8 weeks. Proteomic analysis revealed that the adipose secretome from animals on a high fat diet was enriched in pathways related to immune cell-mediated inflammation, with interleukin 6 (IL-6) as a central regulator of differentially expressed proteins. A multiplex assay to measure cytokines confirmed higher IL-6 in the adipose secretome of high fat-fed animals. Exposure of APCs to IL-6 increased adipogenesis, while treatment of APCs with an IL-6 blocking antibodies diminished the adipogenic effect of the adipose secretome. Together, these findings substantiate a role for redox signaling in the regulation of adipogenesis and identify IL-6 as a novel activator of adipogenesis that may mediate APC differentiation via generation of ROS under obesogenic conditions.

## INTRODUCTION

Adipogenesis is the two-step process by which mesenchymal adipose-derived progenitor cells (APCs) commit to the adipocyte lineage and subsequently undergo terminal differentiation to become lipid-storing adipocytes. High rates of adipogenesis during fetal and early postnatal periods of development coincide with rapid accumulation of fat mass, establishing the setpoint of early life adiposity. APCs residing in mature adipose depots are largely quiescent, as adipocyte number remains relatively stable through adulthood; however, their recruitment for differentiation supports normal adipocyte turnover, or increases capacity for lipid storage under obesogenic conditions (1–3)

The transcriptional regulation of adipogenesis has been characterized in cell lines of embryonic fibroblasts or freshly isolated APCs. Differentiation of multipotent APCs into committed preadipocytes is directed by bone morphogenetic protein (BMP) and Wnt signaling pathways. Upon exposure of growth arrested preadipocytes to adipogenic stimuli, cyclic AMP response element-binding protein (CREB) becomes phosphorylated and activates the expression of CCAAT enhancer binding protein (C/EBP) β which pushes the preadipocyte towards terminal differentiation by inducing transcription of peroxisome proliferator-activated receptor (PPAR)γ, the master regulator of adipocyte genes (4).

While the transcriptional program that drives adipogenesis *in vitro* is well-established, there is a lack of knowledge with respect to the cues that activate APC differentiation under obesogenic conditions. Accumulating evidence suggests that APCs are released from the quiescent state to replace lost adipocytes that have undergone cell death after reaching their expansion capacity (5). Adipocyte death increases during the early stages of obesity, triggering the recruitment of adipose tissue macrophages that adopt a phenotype equipped for lysosome biogenesis and lipid metabolism for phagocytosis of lipid remnants and other cellular debris (6–9). These sites of adipocyte death are located in close proximity to newly generated adipocytes, suggesting that adipogenesis is coupled to adipocyte death in the obese state (5). Thus, a pro-inflammatory signal released during immune cell-mediated adipose tissue remodeling may be the stimulus that triggers APC differentiation.

In our previous work, we identified reactive oxygen species (ROS) as key mediators of the pro-adipogenic effect of endocrine disrupting chemicals in primary culture of murine APCs (10). At physiological levels, ROS function as important second messengers in a variety of signal transduction pathways (11), including cellular responses to inflammatory mediators. Herein, we sought to elucidate the role of redox signaling in the adipogenic behaviour of APCs. Our findings show that activation of transcriptional pathways that drive adipogenesis are accompanied by changes in the expression and activity of antioxidant defenses, and that superoxide (O^2-^) facilitates early differentiation, while hydrogen peroxide (H_2_O_2_) is involved in the lipid accumulation that occurs in later stages of preadipocyte maturation. Further, we reveal IL-6 to be central to the pro-inflammatory environment of the adipose secretome under conditions of obesity and a key mediator of APC differentiation via activation of redox signaling.

## MATERIALS & METHODS

### Animals

All experiments involving animals were conducted at the University of Calgary, approved by the University of Calgary Animal Care Committee (AC21-0132), and conducted in accordance with guidelines by the Canadian Council on Animal Care Ethics. All experiments were conducted using C57BL/6J mice housed in a facility with a 12/12-hour light/dark cycle at 22.5°C and 35% humidity. Animals were humanely euthanized by anesthetic induction under 2% isoflurane followed by decapitation. For *in vitro* culture experiments, adipose tissue was collected from 6– 9-week-old male mice. Male and female mice used in adipose explant experiments were fed a high-fat/high-fructose diet (HFFD, 45% kcal from fat; 35% kcal from fructose, Research Diets Inc., D08040105I) or control diet (CD, 10% kcal from fat, Research Diets Inc., D12450Ki) for 8 weeks beginning at 6 weeks of age.

### Primary APC Isolation and Culture

The inguinal subcutaneous adipose tissue from male mice was dissected under sterile conditions and digested in a 1mg/ml Collagenase I (Worthington Biochemical) digestion buffer in HBSS with 100 mM HEPES and 1.5% BSA for 1 hour at 37°C. The stromal vascular fraction was isolated by centrifugation, followed by incubation in a red blood cell lysis buffer for culture of APCs as previously described (10). Cells were grown in preadipocyte growth media (Cell Applications) until 48 hours past 90% confluency, at which time the media was replaced with adipocyte differentiation media (Cell Applications). After 5 days of culture in differentiation media, differentiated cells were induced to accumulate lipid by the addition of adipocyte maintenance media (Cell Applications) for a further 2 days.

### Explant Culture and Conditioned Media Collection

Inguinal subcutaneous adipose tissue or gonadal visceral adipose tissue was isolated from male and female mice fed a high fat/fructose diet (HFD, Research Diets) or control diet (CD, Research Diets) for 8 weeks. Adipose tissue was aseptically dissected into multiple 10mm pieces and equilibrated in explant media (M199, 1% FBS, 1% P/S) for 24 hours. After 24 hours, explants were rinsed with PBS and the media replaced. After an additional 24 hours, conditioned media was collected and the protein ‘secretome’ isolated using protein concentrators (Pierce). Protein concentrated conditioned media (CM) was frozen at −80°C until use. APCs were treated with CM [500ug/mL] over the course of culture as described above. APCs cultured in the presence of CM were induced to differentiate using low potent differentiation media (High glucose DMEM, 10% FBS, 1% Penstrep, 172nM bovine insulin, 1 uM dexathemasone, 0.5 mM IBMX).

### Manipulation of redox and inflammatory signaling in APCs during differentiation

APCs were treated with the ROS generators: 2-Methylnaphthalene-1,4-dione (Menadione, Sigma-Aldrich) and 2,3-dimethoxy-1,4-napthalenedione (DMNQ, Sigma-Aldrich); the ROS scavengers and antioxidants: N-acetyl cysteine (NAC, Cayman), 4-hydroxy-2,2,6,6-tetramethylpiperidine-N-oxyl (Tempol, Millipore), and mitoquionone (MQ, Cayman), over the course of differentiation beginning at 90% confluency. DMSO (Sigma Life Sciences) was used as vehicle for all treatments except NAC, which is soluble in water. APCs were also treated with the antioxidant catalase (CAT, Sigma Aldrich) and the CAT inhibitor 3-Amino-1,2,4-triazole (3-AT, SigmaAldrich). NFκB inhibition was achieved with diethyldithiocarbamic acid sodium salt trihydrate (DETC, Santa Cruz Biotechnology). APCs were treated with the pro-inflammatory cytokines interleukin 6 protein (IL-6, Stem Cell technologies), or tumour necrosis factor alpha (TNFα, R&D Systems); whereas these pathways were inhibited by treating APCs with IL-6 or TNFα blocking antibodies (Invitrogen) with IgG1κ isotype used as a control (Invitrogen).

### Quantification of APC Differentiation

Differentiated APCs were fixed in 4% PFA for 1 hour at room temperature and lipid droplets were stained with Oil Red O (Sigma-Aldrich). APCs were lysed with lysis buffer (1:25 IGEPAL in 100% isopropanol) for 10 minutes and the absorbance of the eluted dye quantified in triplicates at 490nm on a spectrophotometer (Biotek 800TS). A subset of APCs was fixed for 30 minutes in 4% PFA and stained with 2 μM BODIPY (Thermo Fisher) and 20 μM Hoechst (Thermo Fisher). Images were captured using a Nikon Eclipse Ts2 microscope and processed with NIS-Elements D 5.11.00. Total RNA was isolated using the RNeasy Mini Kit (Qiagen) according to the manufacturer’s instructions, assessed for integrity using an RNA Agilent 4200 TapeStation (University of Calgary Analytical Services) and quantified on an N50 Nanophotometer (Implen Inc.). Following a DNA denaturation step, the High-Capacity cDNA Reverse Transcription kit (Applied Biosystems) was used for cDNA synthesis. cDNA products were mixed in Powerup SYBR green master mix (Applied Biosystems) along with primer pairs and run in triplicates for quantitative real time PCR on a QuantStudio 5 Real-time PCR System (Applied Biosystems). Primers were designed using the NCBI/Primer Blast tool (Supplementary Table 1). Target mRNA expression was normalized to β-actin and calculated using 2^−ΔΔCt^ method, expressed as fold change when comparing vehicle to experimental conditions.

### Quantification of ROS, Antioxidant Activity and Mitochondrial Morphology

APCs were treated with an adipogenic stimulus (differentiation media) or CM for 48 hours for analysis of ROS. Cellular ROS levels quantified using H2DCFDA (Invitrogen) were performed by the fluorescent plate reader method or flow cytometry. APCs were stained with DCFDA in HBSS for 45 minutes at 37°C. For the fluorescent assay, APCs were incubated in DCFDA working solution [10uM] for 45 minutes at 37°C. Fluorescence was quantified at 485nm/535nm using the SpectraMax microplate reader. For flow cytometry, cells were trypsinized (0.25% Trypsin-EDTA 1X, Gibco) and resuspended in cell buffer (HBSS with 5% FBS). The cell suspension was stained with DCFDA working solution [10uM] for 30 minutes on ice. Following incubation and washing, DCFDA positive cells were measured using flow cytometry on the Cytek Aurora Flow Cytometer with unstained cells used as a gating control. Levels of H_2_O_2_ were quantified using the Amplex™ Red Hydrogen Peroxide/Peroxidase Assay Kit (Invitrogen) following the manufacturer’s instructions and fluorescent intensity detected using the SpectraMax microplate reader. Over the course of differentiation, superoxide dismutase (SOD) and CAT activity were determined using the Superoxide Dismutase Assay Kit (Cayman) and Catalase Assay Kit (Cayman), according to the manufacturer’s instructions from samples collected on day 0, 2, 4, and 7 of differentiation or samples treated with CM until D7. APCs acutely (24 and 48 hours) exposed to an adipogenic stimulus or controls in growth media were stained with Mitotracker deep red (Thermo Fisher) at a 1:1000 dilution and incubated for 1 hour at 37°C. Live cells were captured on a Ti2 ZDrive Nikon A1plus at 60x and Z-stacks captured with 0.2 μ steps. In every image, mitochondria of each cell were assigned as elongated (E), intermediate (I), or shortened (S). The proportion of E, I, and S cells was determined for each image and averaged for two blinded evaluators.

### Proteomic Analysis of the Adipose Secretome

Shotgun proteomics were performed on concentrated protein from CM. Total protein concentrations were determined using Pierce BCA Assay kit according to the manufacturer instructions. Samples were prepared using the filter-assisted separation of peptides (FASP) method for which 100μ of protein was precipitated by the addition of trichloroacetic acid (TCA). Samples were centrifuged at 14,000 x *g* for 15 minutes at 4°C. Subsequently, cells were washed 3 times in cold acetone and stored at −20°C overnight, then resuspended in 8M tris-urea solution by shaking and denatured with the addition of 10 mM DTT at 37°C for 30 minutes. To complete the carbamidomethyl modification of cystines, 50 mM of iodoacetamide was added in the dark at room temperature, then samples were centrifuged at 14,000 x *g* on top of a 30 kDa filter for 15 minutes. Samples were washed with 8M tris-urea, followed by 50mM ammonium bicarbonate. The samples were trypsinized at 37°C overnight at a ratio of 1:10 trypsin: total protein, eluted off the filter membrane by washing with 50mM ammonium bicarbonate, then subjected to a c18 clean up with waters solid-phase extraction (SPE) column, according to directions manufacturer instructions (Waters).

### High performance liquid chromatography (HPLC) and mass spectrometry (MS)

Using a protocol previously described by Mishra et al. (12), tryptic peptides were analyzed on an Orbitrap Fusion Lumos Tribrid mass spectrometer (Thermo Scientific) operated with Xcalibur (version 4.4.16.14) and coupled to a Thermo Scientific Easy-nLC (nanoflow Liquid Chromatography) 1200 system. Tryptic peptides (2 μ were loaded onto a C18 trap (75 um x 2 cm; Acclaim PepMap 100, P/N 164946; Thermo Scientific) at a flow rate of 2μl/min of solvent A (0.1% formic acid in LC-MS grade water). To elute the peptides, a 120 min gradient from 5 to 40% (5% to 28% in 105 min followed by an increase to 40% B in 15 min) of solvent B (0.1% formic acid in 80% LC-MS grade acetonitrile) at a flow rate of 0.3 μL/min and separated on a C18 analytical column (75 um x 50 cm; PepMap RSLC C18; P/N ES803; Thermo Scientific). Peptides were then electrosprayed using 2.1 kV voltage into the ion transfer tube (300°C) of the Orbitrap Lumos operating in positive mode. The Orbitrap first performed a full MS scan at a resolution of 120,000 FWHM to detect the precursor ion having a m/z between 380 and 985 and a +2 to +7 charge. The Orbitrap AGC (Auto Gain Control) and the maximum injection time were set at standard and 50 ms, respectively. The Orbitrap top speed mode with a 3 sec cycle time for precursor selection was used. The most intense precursor ions presenting a peptidic isotopic profile and having an intensity threshold of at least 5,000 were isolated using the quadrupole and fragmented with HCD (30% collision energy) in the ion routing multipole. The fragment ions (MS2) were analyzed in the orbitrap with a 40 sliding windows of 16.0072 m/z. The AGC and the maximum injection time were set at 1×10^4^ and 35 ms, respectively, for the quadrupole.

### Bioinformatic Analysis

Spectra data obtained during mass-spectrometry were matched to peptide sequences from a Mouse FASTA reference file obtained from Uniprot on Jan 11^th^ 2024 using DIA-NN (v.1.8.1). DIANN was run in “FASTA digest for library-free search mode” Variable modifications included N-term M excisions. Fixed modifications included Carbamidomethylation on Cystines. All other DIANN settings were set to default. Then MSstatsShiny UI (v0.1.0; https://msstatsshiny.com/app/MSstatsShiny) running MSstats (v.4.2.0) was used to run the statistical analysis to compare CD male subcutaneous CM samples to HFD male subcutaneous CM samples. The DIANN file “Report.tsv” were used for the analysis, while the annotation file was created according to directions provided by MSstats. Unique peptides were included in the analysis. For data processing, log2 was defined for data transformation and equalize medians was the method chosen for normalization. The options “Use unique peptides”, “remove proteins with 1 feature”, and “remove runs with over 50% missing values” were used for this analysis. Statistical inference was performed using custom pairwise comparisons, and the data was imported to excel for final assessment.

### Protein–Protein Interactions and Pathway Analysis

To investigate the biological significance of differentially expressed proteins, Ingenuity Pathway Analysis (IPA) software (Qiagen Inc.) was used to further analyze significantly enriched or decreased (p<0.1) proteins. IPA is a widely used tool that can be used to predict biological outcomes of expression differences. Datasets containing protein identifiers (UniProt) and corresponding expression values (Log2 [Fold change]) of male CD vs HFD were uploaded, and predicted networks were analyzed.

### Cytokine Quantification

The secretome collected from CM of male CD and HFD subcutaneous explants were submitted to Eve Technologies Corp. (Calgary, Alberta) for Luminex multiplex ‘Mouse Cytokine Proinflammatory Focused 10-Plex Discovery Assay® Array (MDF10) to quantify cytokine expression [GM-CSF, IFNγ, IL-1β, IL-2, IL-4, IL-6, IL-10, IL-12p70, MCP-1, TNFα]. The multiplexing analysis was performed using the Luminex™ 200 system (Luminex, Austin, TX, USA). The markers were simultaneously measured in the samples using Eve Technologies’ Mouse Focused 10-Plex Discovery Assay® (MilliporeSigma, Burlington, Massachusetts, USA) according to the manufacturer’s protocol. Assay sensitivities of these markers range from 0.4 – 10.9 pg/mL for the 10-plex. Individual analyte sensitivity values are available in the MilliporeSigma MILLIPLEX® MAP protocol.

### Statistical Analysis

Comparisons between two groups were made using an unpaired student’s t-test. Analyses with multiple groups were performed using a one-way ANOVA followed by Dunnett’s *post hoc* test. Comparisons across depot and diet for CM experiments were compared using a two-way ANOVA followed by uncorrected Fischer’s least significant differences test. Graphs are represented as mean ± SEM unless otherwise stated.

## RESULTS

### APC differentiation coincides with increased ROS production and is impacted by ROS levels

First, we set out to determine the impact of APC differentiation on intracellular levels of ROS. Compared to the undifferentiated state, APCs that were exposed to differentiation media for 48 hours exhibited increased levels of O_2_^-^, as measured by DCFDA (Figure 1A-B). Short-term exposure of APCs to differentiation media did not impact levels of H_2_O_2_, measured by Amplex Red staining (Figure 1C). Since O_2_^-^ levels were found to increase during the early stages of adipogenesis, ROS levels were pharmacologically manipulated to determine the impact on adipogenic capacity of APCs. Exposure of APCs to redox cyclers, DMNQ and menadione, which stimulate the production of O_2_^-^ and H_2_O_2_, augmented the differentiation of APCs, as reflected by increased lipid droplet staining (Fig 1D-G). Over the course of differentiation, the peak in mRNA expression of PPARγ, the master regulator of APC differentiation, occurred earlier in APCs exposed to menadione and remained elevated during the maintenance period, whereas expression levels declined after day 5 in APCs cultured in vehicle control. The peak in mRNA expression levels of glucose transporter type 4 (GLUT4), a marker of differentiated APCs, occurred on day 6 of differentiation, and was amplified in APCs exposed to menadione (Fig 1F). APCs treated with NAC, which has both ROS scavenging and antioxidant properties and broadly decreases intracellular ROS, exhibited blunted adipogenic potential with lower lipid droplet formation on day 4 and day 7 of differentiation (Fig 1H-I). Reduction of ROS via NAC was found to diminish the expression of PPARγ and GLUT4 over the course of differentiation compared to vehicle (Fig 1J). Treatment with tempol, which prevents iron oxidation and thereby broadly reducing ROS production, decreased lipid droplet staining on day 7 of differentiation (Fig 1K). To specifically target mitochondria-derived ROS, APCs were treated with MQ, a mitochondrial-targeted antioxidant, which decreased lipid droplet staining on day 7 only (Fig 1L). Mitochondrial length was impacted by differentiation with more short mitochondria in differentiating cells (Supplementary Fig 1).

**Figure 1.**
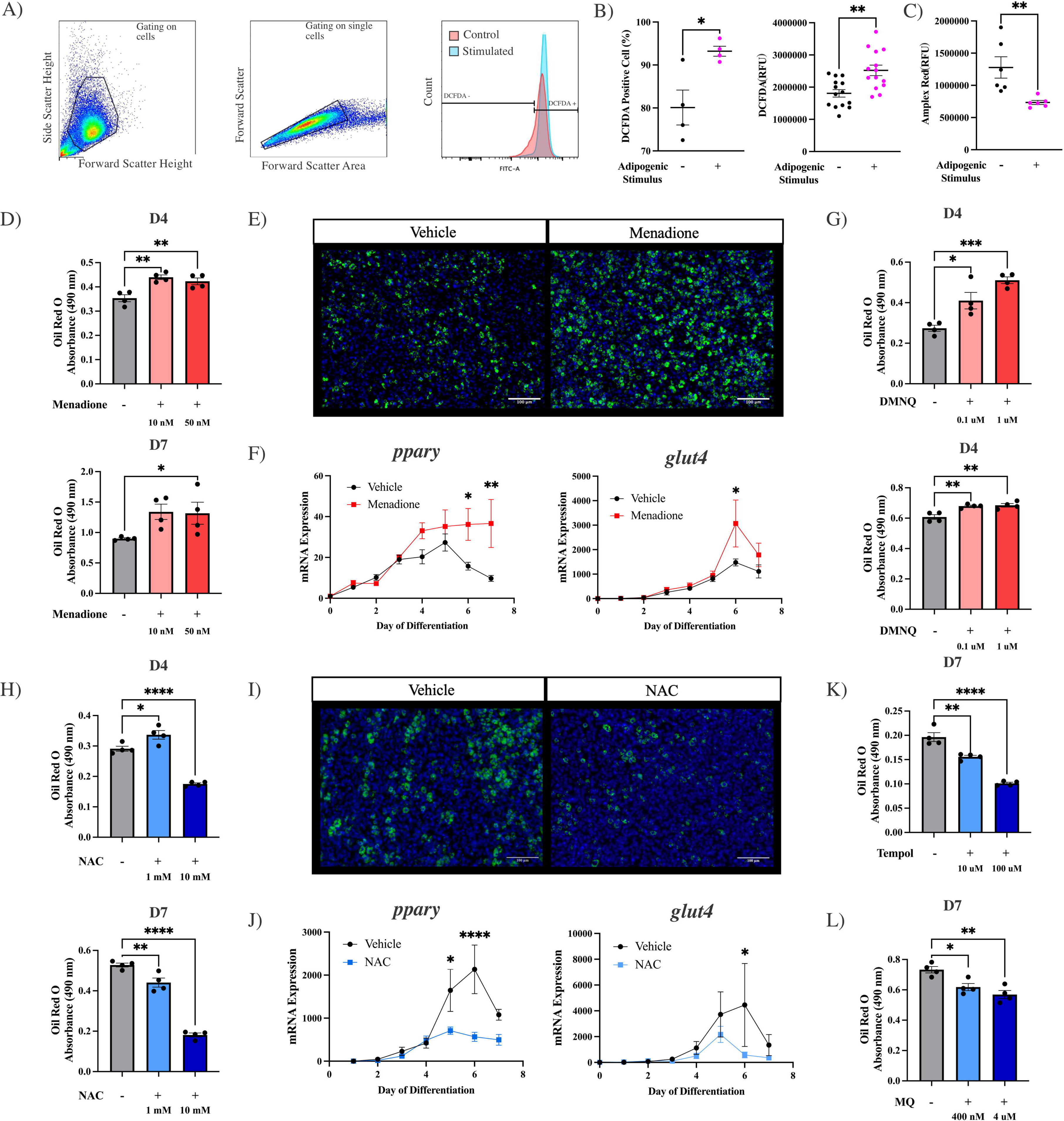
ROS Increases Adipogenesis in Primary APC Culture. (A) Representative plots of flow cytometric identification of DCFDA-positive APCs. (B) APCs cultured in adipogenic media for 48 hours were stained with DCFDA, which was quantified by flow cytometry or by the microplate reader method. (C) APCs were cultured in adipogenic media for 48 hours and stained with Amplex Red, for the measurement of H_2_O_2_. (D) Absorbance of eluted Oil Red O from stained lipid droplets of APCs treated with ROS-generator, DMNQ, or vehicle on day 4 and day 7 of differentiation. (E) APCs stained with BODIPY (lipid droplet) and Hoescht (nuclei) on day 7, with or without exposure to ROS-generator, menadione. (F) Fold change relative to vehicle of mRNA expression of *pparγ* and *glut4* from day 0 to day 7 of differentiation in APCs treated with ROS-generator, menadione [50nM]. (G) Eluted Oil Red O absorbance from stained APCs exposed to menadione or vehicle and collected on day 4 and day 7 of differentiation. (H) Eluted Oil Red O absorbance from APCs treated with NAC or vehicle and collected on day 4 and day 7 of differentiation. (I) BODIPY and Hoescht-stained APCs on day 7 of differentiation in the presence or absence of the antioxidant, NAC. (J) Fold change relative to vehicle of mRNA expression of *pparγ* and *glut4* from day 0 to day 7 of differentiation in APCs treated with NAC [10 mM] or vehicle. (K) Eluted Oil Red O absorbance from APCs treated with the ROS scavenger, Tempol, and collected on day 7 of differentiation. (L) Eluted Oil Red O absorbance from APCs treated with the mitochondria-targeted antioxidant, MQ, and collected on day 7 of differentiation. * p ≤ 0.05, ** p ≤ 0.01; *** p ≤ 0.001, **** p ≤ 0.0001.

### APC Differentiation Corresponds with Changes in the Expression and Activity of Antioxidants

Given our findings that ROS levels change over the course of adipogenesis, and manipulation of ROS can modulate the degree of differentiation in response to an adipogenic stimulus, we determined whether changes in antioxidant gene expression and activity parallel adoption of the adipogenic transcriptional program during adipogenesis. The antioxidant SOD is the major cellular response against O^2-^ and exists in three isoforms. SOD1, the cytoplasmic isoform, significantly increased over the course of differentiation, starting at day 0 upon initiating differentiation and peaking at day 5, with a decline thereafter (Fig. 2A). SOD2, the SOD isoform present in mitochondria, exhibited a biphasic expression pattern, peaking early with a sharp increase from day 0 to day 1 and a second lower peak on day 5 (Fig. 2A). Similarly, the extracellular isoform, SOD3, peaked on the first day after initiation of differentiation, although changes in SOD3 over the course of differentiation were relatively subtle (Fig 2A). The mRNA expression of glutathione peroxidases (GPX), which catalyzes the reduction of H_2_O_2_, increased from day 3 of differentiation peaking on day 6 (Fig 2A). Similarly, the mRNA expression of the H_2_O_2_ targeted antioxidant, CAT, increased throughout differentiation, with a peak on day 5 and a decline thereafter (Fig 2A). We then quantified the overall activity of SOD over the course of adipogenesis and found activity to increase from day 2 onward with a peak at D4 (Fig 2B). The enzymatic activity of CAT was increased on day 4 of differentiation (Fig 2C).

**Figure 2.**
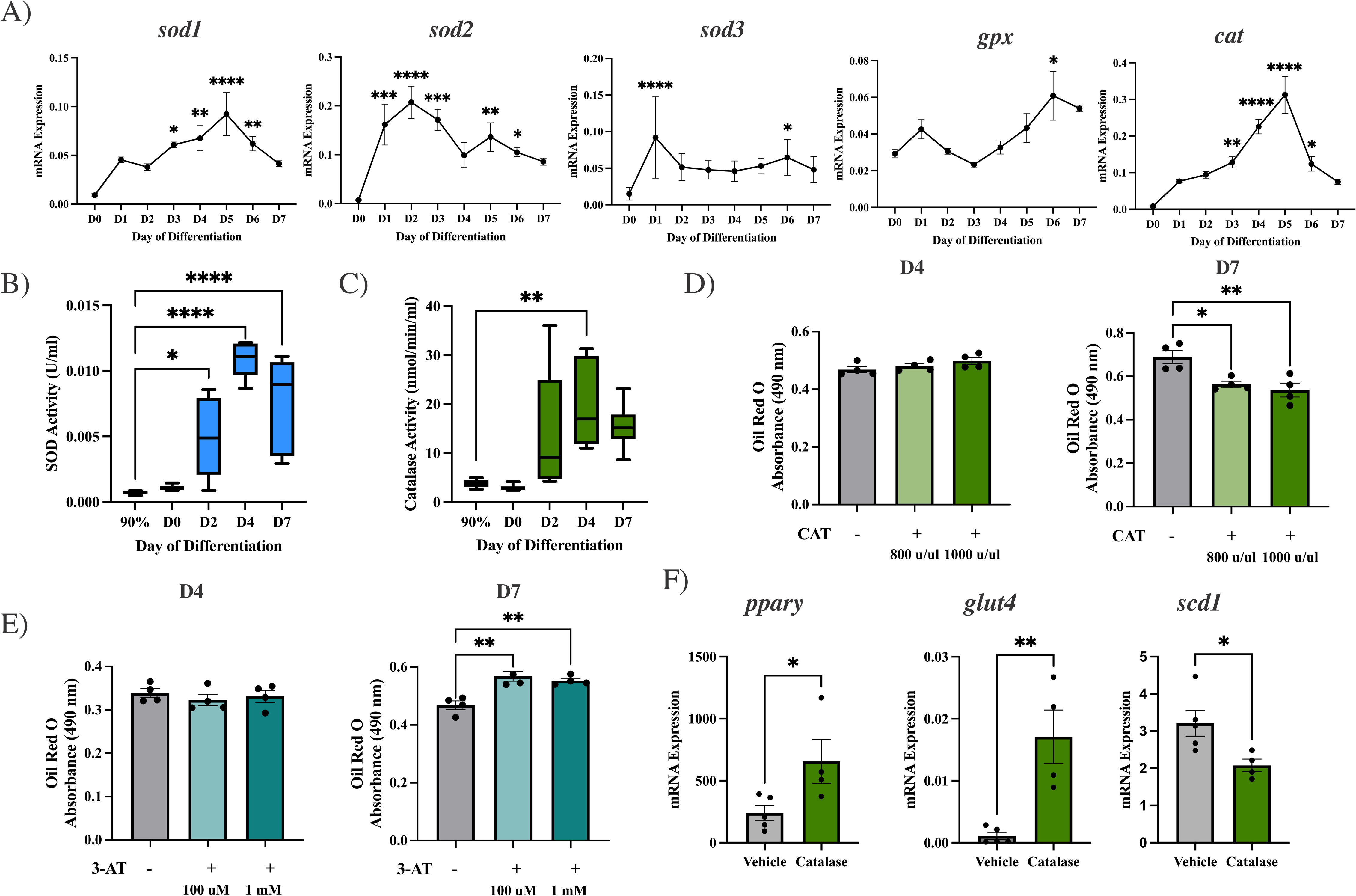
Antioxidant Defenses are Activated During Adipogenesis. (A) Fold change relative to vehicle of mRNA expression of SOD isoforms *sod1*, *sod2*, and *sod3, glutathione peroxidase* (*gpx*), and catalse (*cat*) from day 0 to day 7 of differentiation (n=6). (B) SOD activity from 90% confluency (48 hours before day 0) to day 7 of differentiation (n=6). (C) CAT activity from 90% confluency (48 hours before day 0) to day 7 of differentiation (n=6). (D) Eluted Oil Red O absorbance from stained lipid droplets of APCs treated with CAT or vehicle and collected on day 4 and day 7 of differentiation (n=4). (E) Eluted Oil Red O absorbance from APCs treated with 3-AT (Cat inhibitor) or vehicle and collected on day 4 and day 7 of differentiation (n=4). (F) mRNA expression of adipogenic genes *pparγ* and glut4, and *scd1*, involved in lipid synthesis, in APCs treated with CAT or vehicle and collected on day 7 of differentiation (n=4-5). * p ≤ 0.05, ** p ≤ 0.01; *** p ≤ 0.001, **** p ≤ 0.0001.

### Hydrogen peroxide influences lipid droplet formation in late adipogenesis

Temporal differences in the mRNA expression and activity of antioxidant defenses against O_2_^-^ vs. H_2_O_2_ suggest that these radical species play different roles during APC differentiation. Since H_2_O_2_ is a byproduct of the dismutation of O_2_^-^ by SOD and increased along with higher O_2_^-^ production, we isolated the impact of H_2_O_2_ by treating APCs with CAT. Treatment of APCs with CAT, which reduces H_2_O_2_, decreased lipid staining on day 7 only with no changes found on day 4 (Fig 2D). Conversely, increasing H_2_O_2_ levels by treating APCs with a CAT inhibitor (3-AT), increased lipid droplet staining on day 7 only (Fig 2E). After switching differentiation media for maintenance on day 5 of differentiation, lipid droplets accumulate in differentiated APCs. Thus, the influence of H_2_O_2_ may be restricted to lipogenesis in late adipogenesis, whereas O_2_^-^ appears to impact early differentiation. Indeed, reducing H_2_O_2_ levels with CAT decreased mRNA expression of Stearoyl CoA desaturase 1 (SCD1), a key enzyme in lipogenesis, but increased levels of PPARγ and GLUT4 (Fig. 2F), providing further evidence that H_2_O_2_ facilitates lipid droplet formation in differentiated APCs rather than stimulating early APC differentiation.

### The obese adipose secretome contains signals that activate adipogenesis via redox signaling

Given that redox signaling orchestrates cellular responses to pro-inflammatory signals, and release of pro-inflammatory adipokines such as TNFα, IL-6 and IL1β occurs in obesity, we set out to determine if redox signaling mediates the activation of APC differentiation in response to signals present in the adipose secretome. First, to establish that adipokines released in the obese state activate APC differentiation, we exposed cultured APCs to the adipose secretome. Adipose explants were isolated and cultured *ex vivo* for the collection of CM, from which the secretome was extracted by concentrating proteins, thereby eliminating the influence of lipids (Fig 3A). In the presence of CD or HFD male CM from subcutaneous explants, adipogenesis was enhanced compared to APCs differentiated in the absence of CM (Supplementary Fig. 2), suggesting that secreted factors from adipose tissue exhibit pro-adipogenic properties. We then compared CM from male mice fed a CD vs HFD for 8 weeks, which temporally corresponds to the activation of the *in vivo* adipogenic response in high fat diet-fed mice (2). APCs differentiated in the presence of CM from visceral adipose explants of HFD-fed male mice exhibited higher lipid droplet staining on day 4 and day 7; whereas lipid droplet staining was higher on day 7 only in APCs differentiated in the presence of CM from subcutaneous adipose explants of HFD-fed male mice. (Fig 3B). A similar pattern was exhibited following treatment with CM isolated from female adipose tissue, demonstrating enhanced adipogenesis on day 4 only in the presence of HFD-CM from subcutaneous depots and on day 7 only in the presence of HFD-CM from visceral depots (Fig 3B). Overall, HFD-CM has a greater pro-adipogenic effect than CM from CD-fed animals, suggesting that pro-adipogenic adipokines are increased in the early stages of obesity when adipogenesis is activated in *in vivo* models.

**Figure 3.**
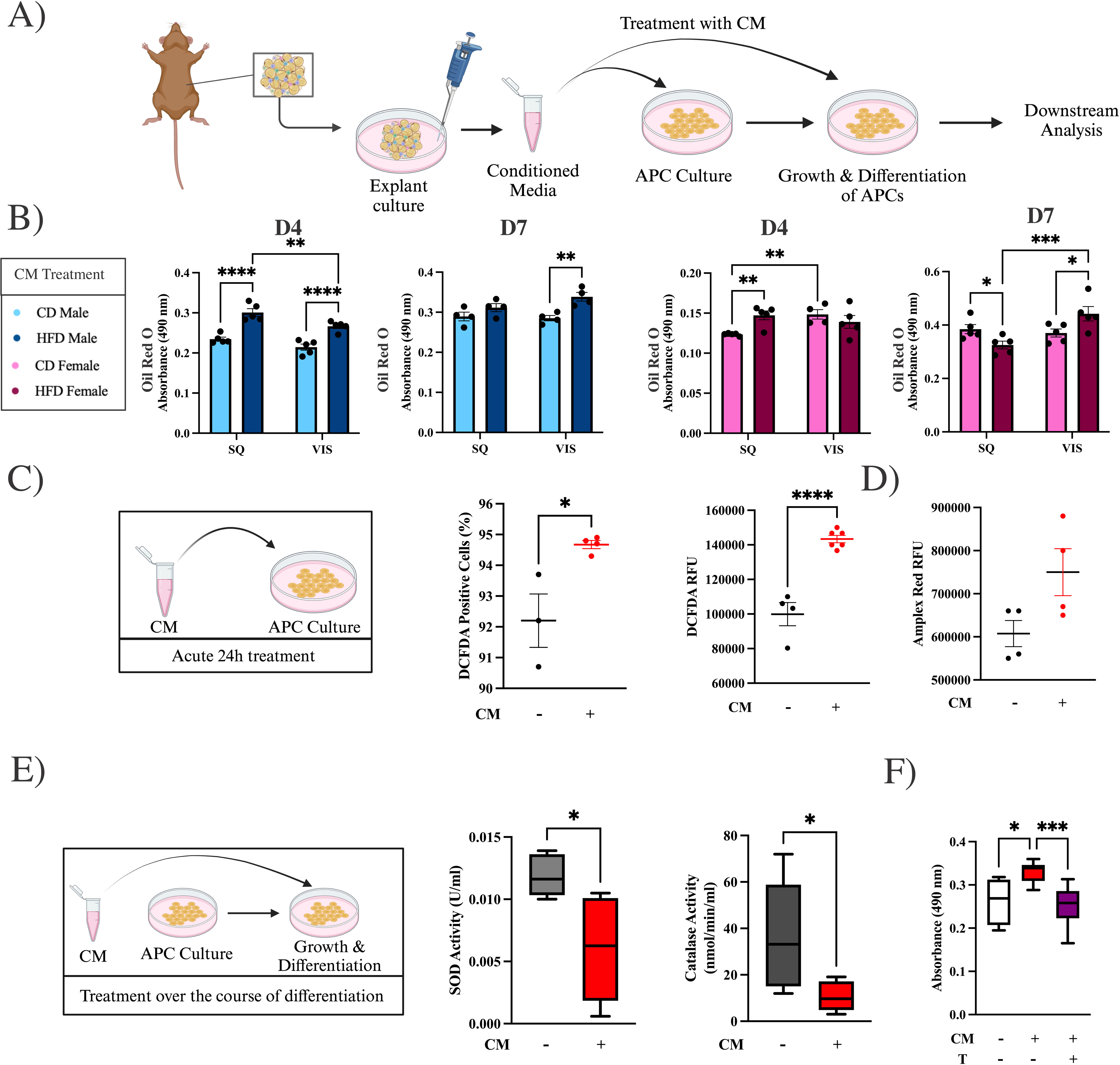
The Secretome of Adipose Explants Enhances Adipogenesis and ROS Production Under Obesogenic Conditions. (A) Schematic of conditioned media experiments. (B) Eluted Oil Red O absorbance on day 4 and 7 from stained lipid droplets of APCs differentiated in the presence of proteins concentrated from conditioned media (CM) collected from adipose explants of male or female inguinal subcutaneous or gonadal visceral depots after 8 weeks of control diet (CD) or high fat diet (HFD) (n=5). (C) APCs treated acutely with CM and stained with DCFDA to measure ROS, which was quantified by flow cytometry and the plate reader method (n=4-6). (D) APCs treated acutely with CM and stained with Amplex Red for quantification of H_2_O_2_ (n=4-6). (E) Superoxide dismutasae (SOD) and catalase (CAT) activity on D7 of APCs differentiating in the presence of CM (n=6). (F) Eluted Oil Red O absorbance in APCs treated with CM with or without the ROS scavenger, tempol (n=4). * p ≤ 0.05, ** p ≤ 0.01; *** p ≤ 0.001, **** p ≤ 0.0001.

Next, we determined if acute exposure of APCs to CM activated redox signaling. Production of O^2-^ measured by DCFDA staining was higher in APCs exposed to CM (Fig 3C); whereas there was a non-significant trend towards higher levels of H_2_O_2_ measured by Amplex Red (Fig 3D). On day 7 of differentiation, APCs differentiated in the presence of CM had lower activity of SOD and CAT activity (Fig 3E). To elucidate the potential role of ROS in mediating the pro-adipogenic effect of CM, APCs were differentiated in the presence of CM with or without the ROS scavenger, tempol. The increase in lipid droplet staining in APCs differentiated with CM was blunted with scavenging of ROS by tempol (Fig 3F).

### The adipokine, IL-6, is central to enriched pathways in the obese adipose secretome

To identify the pro-adipogenic signal within the adipose secretome, CM samples from subcutaneous adipose explants from CD and HFD-fed male mice were subjected to proteomic analysis (Fig 4A). As shown in the volcano plot, 299 proteins were differentially expressed between CM of control vs. HFD (1 ≤ log2fold change ≤ −1; ≥ p = 0.05) with 98 being upregulated (Fig 4B). Among upregulated proteins were those involved in fatty acid oxidation, including ACOX1, and inflammation including LTF, IL-6, and TNFRSF1B (Fig 4B). Several mitochondrial-related proteins including MRPS23, CISD3, EEF2, and MT-CO3 were downregulated (Fig 4B). Canonical pathway analysis in IPA was used to further delve into functional differences based on differential protein expression between CD and HFD secretomes. This analysis revealed enrichment in pathways involved in the cellular immune response, including pathways predicted to upregulate ROS and nitric oxide secretion from macrophages; pathways involved in cellular growth, proliferation and development; cellular stress and injury, the immune system, extracellular matrix organization and metabolism of proteins, including regulation of insulin-like growth factor (Fig 4C).

**Figure 4.**
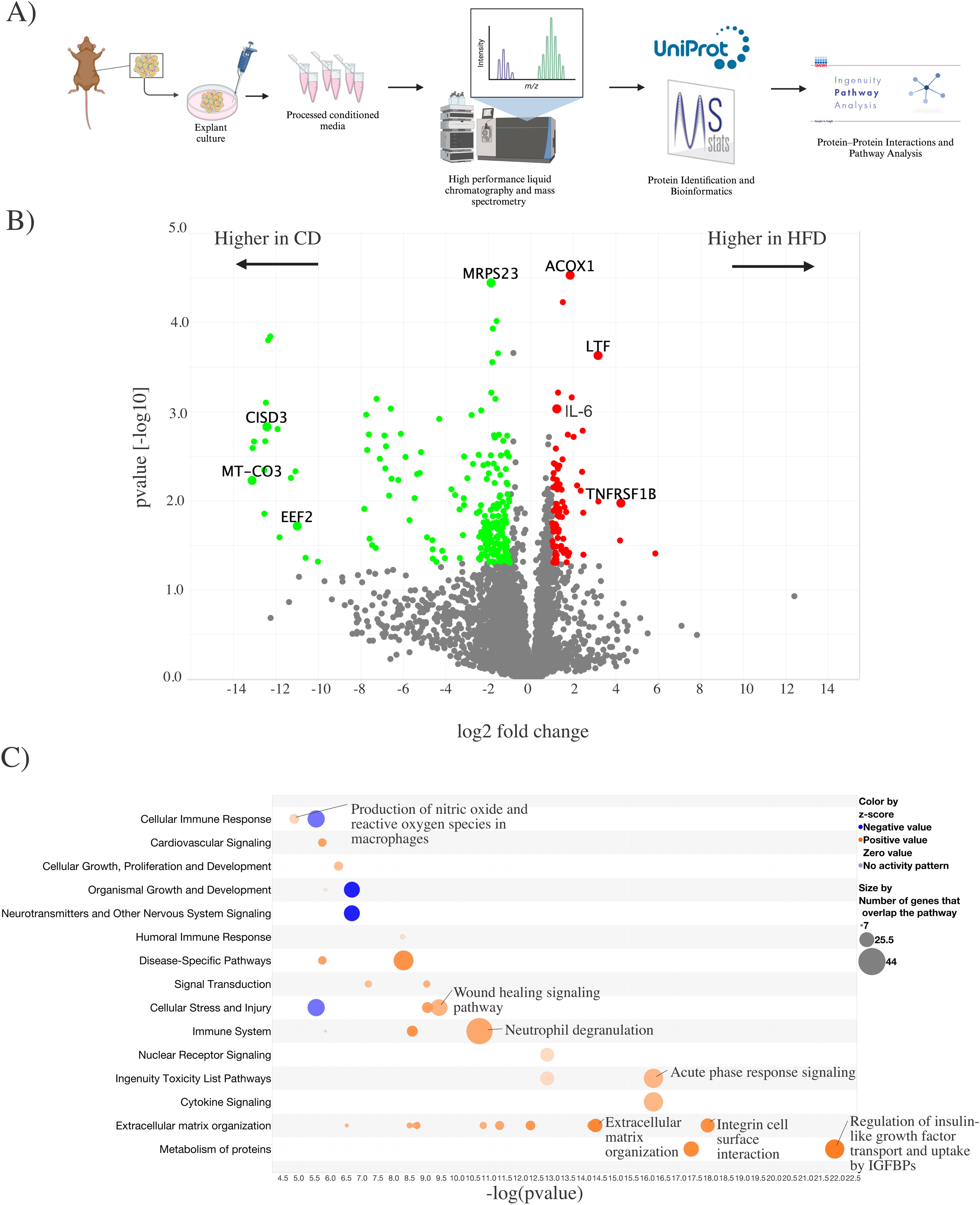
The Protein Secretome of Mice Fed a HFD is Enriched in Pathways Involved in Inflammation and Cellular Growth. (A) Schematic of proteomics pipeline. (B) Volcano plot comparing control diet (CD)-fed male to high fat diet (HFD)-fed males with log2fold change on the x-axis and p value[-log10] on the y-axis, depicting a log2fold change cut off of −1 and 1, and a p value cut off of 0.05 (n=5). (C) Bubble plot displaying differentially regulated pathways identified in IPA canonical signaling.

The most significantly changed proteins were identified by raising the p-value cutoff to 0.01 (Fig 5A). Among activated proteins were IL-6, complement 6 and proteins involved in extracellular matrix remodeling, like MMP3. A graphical summary of the most impactful protein interactions predicted within our dataset reveal IL-6 to be central to the differentially expressed proteins between the adipose secretomes of CD vs. HFD (Fig. 5B). The summary highlights several immune signaling proteins including TNFα, NFκB-1, and IL-1β among others, which were predicted to be activated in our dataset (Fig 5B). Using upstream regulator analysis, IL-6 and TNFα were predicted upstream regulators of many of the differentially expressed proteins in our data set (Supplementary Figure 3).

**Figure 5.**
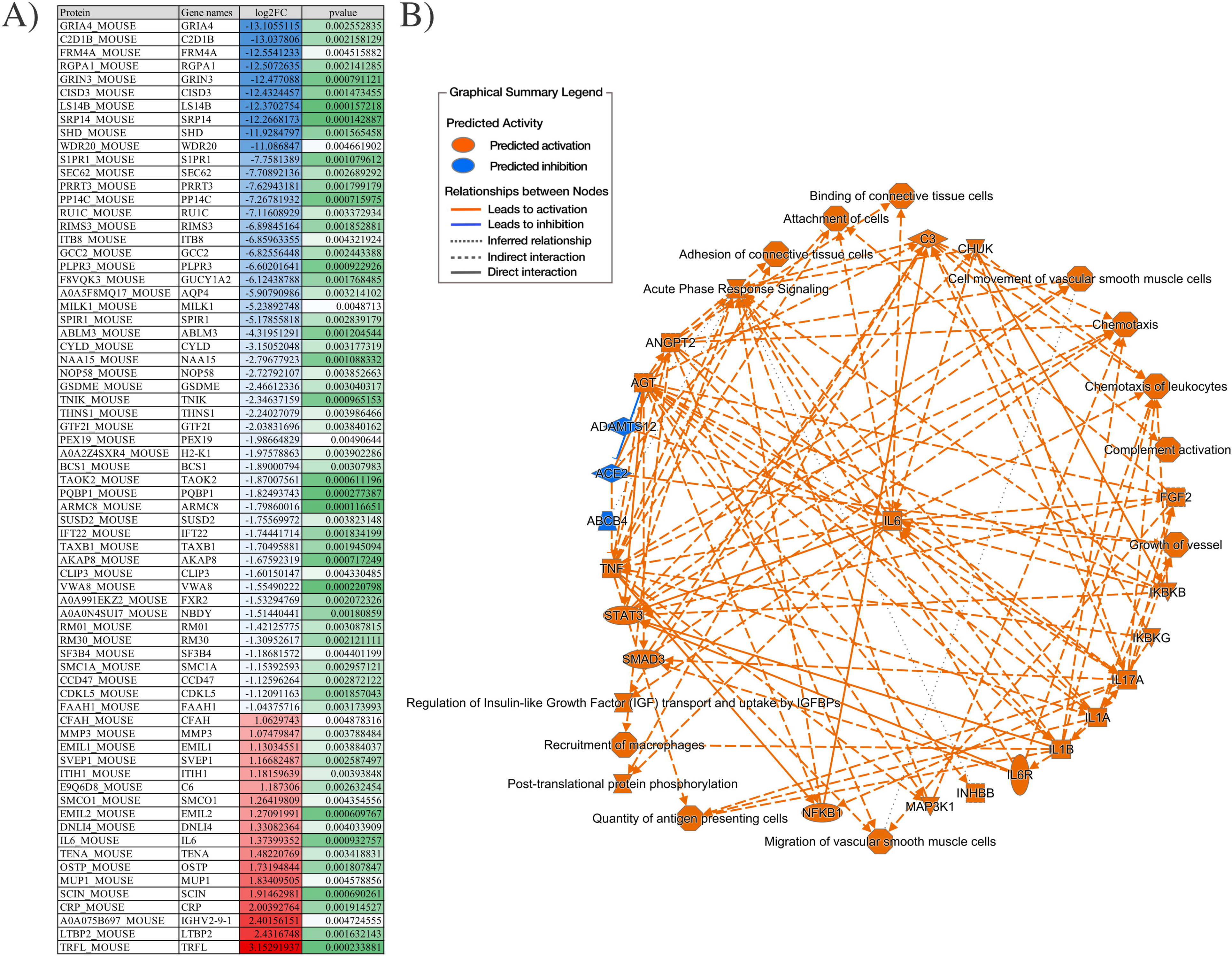
IL6 is Central to the Upregulation of Pro-inflammatory Proteins in the HFD Secretome. (A) Visual representation of the most significantly differentially regulated proteins with a p value cut off of 0.01, depicting protein code, gene name, log2fold change and p value. (B) Visual graphical summary of predicted activation and inhibition of proteins and pathways made in IPA.

### IL-6 mediates the potentiation of adipogenesis in the presence of the adipose secretome

To substantiate differences in cytokine production between control and HFD adipose secretomes, cytokine levels were quantified in CM using a 10-cytokine multiplex assay. There were significant increases in pro-inflammatory cytokines, IL-6 and TNFα, in HFD compared to CD CM samples (Fig 6A). Additionally, the anti-inflammatory cytokine, IL-10, was also upregulated in the HFD group (Fig 6A). There was also a non-significant trend towards increased chemoattractant protein 1 (MCP-1), a chemokine involved in immune cell recruitment. No differences were seen in the expression of the cytokines IL-2 or IL-1β (Fig 6A).

**Figure 6.**
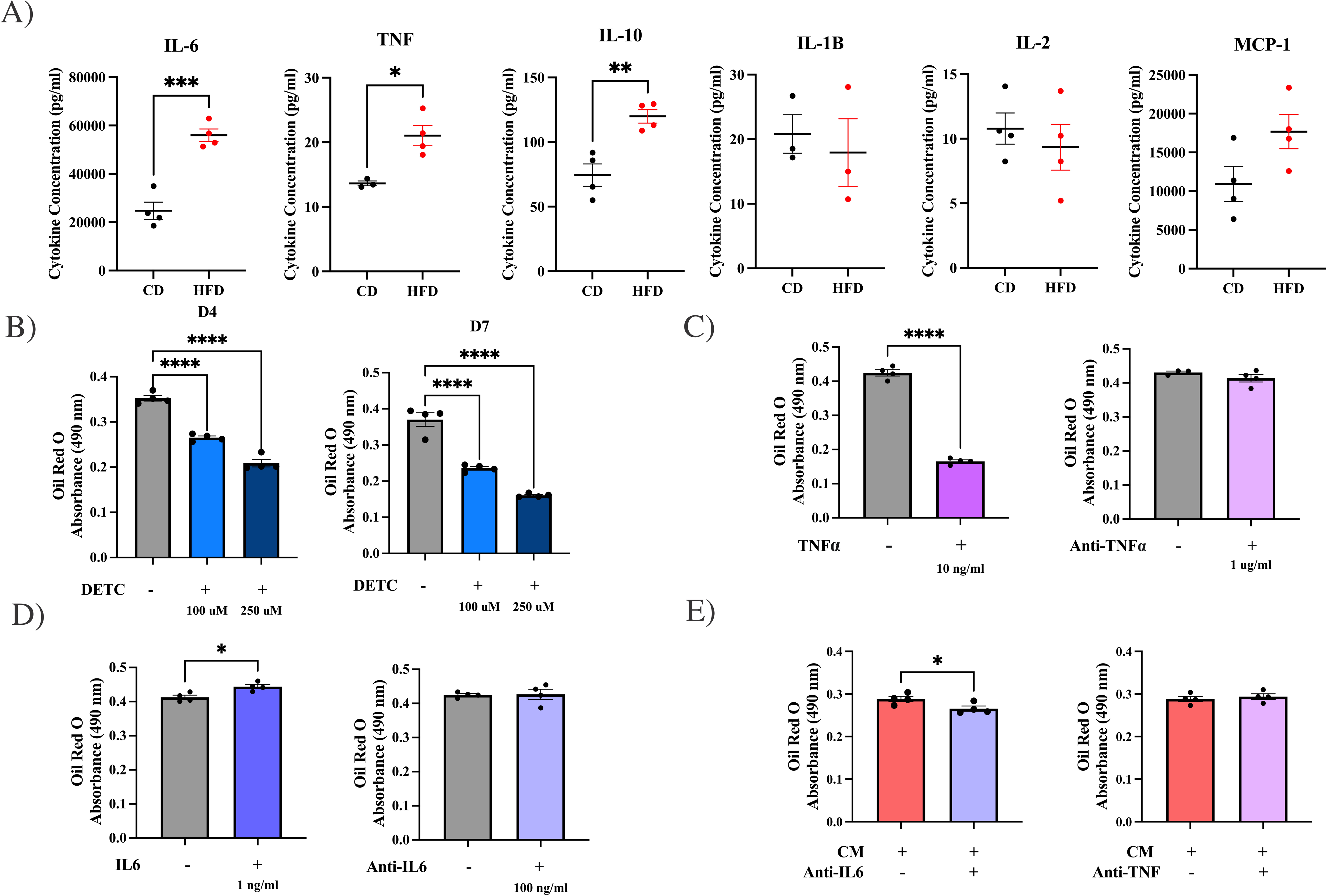
IL6 Derived from the Adipose Secretome Mediate Adipogenesis via Redox Signaling. (A) Concentration of cytokines: IL6, TNFa, IL-10, IL1B, IL-2, and MCP-1 quantified by multiplex assay (n=4). (B) Eluted Oil Red O absorbance from stained lipid droplets of APCs treated with DETC (NFkB inhibitor) and collected on day 4 and 7 of differentiation (n=4). (C) Eluted Oil Red O absorbance from APCs treated with TNFα or TNFα neutralizing antibody and collected on day 7 of differentiation (n=4). (D) Eluted Oil Red O absorbance from APCs treated with IL6 or IL6 neutralizing antibody and collected on day 7 of differentiation (n=4). (E) Eluted Oil Red O absorbance from differentiated APCs co-treated with CM and IL6 or TNFα neutralizing antibodies (n=4) * p ≤ 0.05, ** p ≤ 0.01; *** p ≤ 0.001, **** p ≤ 0.0001. Note: sample size for IL1B is lower due to replicates below detectable threshold.

Next, we sought to determine if pro-inflammatory signals derived from the adipose secretome were involved in adipogenesis. APCs were differentiated in the presence of DETC, an inhibitor of NFκB, a transcription factor central in the crosstalk between inflammatory and redox signaling. Inhibition of NFκB decreased adipogenesis as quantified by lipid droplet staining on D4 and D7 (Fig 6B). Treating APCs with TNFα markedly blunted differentiation, while neutralizing TNFα had no impact (Fig 6C). In contrast, treatment of APCs with IL-6 enhanced APC differentiation, and neutralization of IL-6 did not affect differentiation (Fig. 6D), suggesting that IL-6 may be an activator of adipogenesis but is not required for adipogenesis in the presence of adipogenic media. To determine if IL-6 mediates the augmentation of adipogenesis in the presence of the adipose secretome, APCs were differentiated in the presence of CM with or without IL-6 neutralizing antibodies. Lipid droplet staining was decreased when IL-6 was neutralized during exposure to CM.

## DISCUSSION

The recruitment of APCs for differentiation is a key participant in adipose remodeling. Adaptive adipose remodeling under obesogenic conditions buffers toxic lipids and is thought to distinguish metabolically healthy obesity from metabolically unhealthy obesity (13,14). *In vitro* models for the study of APCs, primarily the 3T3-L1 cell line of embryonic fibroblasts, have determined the transcriptional program of adipogenesis, but have been less useful for understanding the regulation of adipogenesis *in vivo*. Jeffery *et al*. identified that *in vivo* adipogenesis in response to an obesogenic environment is sex- and depot-dependent, and that the adipogenic behaviour of transplanted APCs is governed by the recipient adipose depot rather than the depot of APC origin (15). Therefore, activation of APC differentiation in the obese state is likely driven by paracrine signals arising from neighbouring cells in the local tissue microenvironment. However, the adipogenic signals mediating crosstalk between APCs and other cells in the adipose environment remain unknown.

In our recent work, we discovered that the commonly observed adipogenic effect of endocrine disrupting chemicals is in fact due to an increase in the production of ROS at levels that did not compromise cell viability (10). Interestingly adipogenesis was blunted in cells not exposed to toxicants but treated with ROS scavengers. Our work presented herein further support a role of redox signaling in the activation of APC differentiation and reveal that the transcriptional program of adipogenesis involves upregulation in the expression and activity of antioxidant enzymes. A role for ROS in the regulation of adipogenesis has been demonstrated previously by a handful of studies. Using 3T3-L1 cells, Lee *et al*. found that administration of H_2_O_2_ accelerated adipogenesis (16). Using the 10T1/2 multipotent stem cell line, Kanda *et al*. observed that NAC blocked differentiation and that silencing of NADPH oxidase 4 (Nox4), a generator of H_2_O_2_, abrogated APC differentiation (17). Nox4 has also been implicated in differentiation of human preadipocytes (18). However, Tormos *et al*. found that mitochondrial-targeted antioxidants decreased adipogenesis, similar to results reported herein, but treatment with H_2_O_2_ attenuated this effect (19). Further, this study demonstrated that O_2_^-^ deriving from complex III of the electron transport chain activated adipogenesis independent of oxidative phosphorylation. Han *et al*. found O^2^- generation enhanced in murine APCs isolated from a MnSOD knockout mouse, which increased the number of adipocytes compared to wildtype, but only in HFD fed mice within the gonadal visceral depot (20). An increase in mitochondrial content has been shown to coincide with APC differentiation (19,21). Our results showed that the effects of mitochondria-derived ROS were restricted to late stages of adipogenesis and that mitochondrial length changes over the course of differentiation. In contrast to the above findings, Fernando *et al*. induced oxidative stress in 3T3-L1 cell in hyperoxic conditions and reported a decrease in adipogenesis (22). This highlights the importance of distinguishing dysfunctional from physiological states, the latter wherein ROS act as secondary messengers in normal cellular processes like differentiation. For this reason, we chose to use antioxidants and pharmacological manipulators of redox signaling that are more conductive to titration of ROS levels, rather than genetic approaches, to study the impact of redox signaling on the regulation of adipogenesis.

As described above, while ROS have been demonstrated to influence APC differentiation, the role of specific radical species remains unclear. Our results presented herein suggests that O^2-^ plays a more important role in the early activation of differentiation, whereas H_2_O_2_ appears to be more involved in late adipogenesis at which point lipogenic pathways drive accumulation of lipid droplets in newly differentiated adipocytes. Further investigation is required to substantiate the differential roles of these two radical species, and elucidate the sources of ROS that may include NADPH oxidases and other ROS-producing enzymes or mitochondria.

Since obesity is well known to be associated with a pro-inflammatory adipose environment and redox signaling orchestrates cellular responses to inflammatory signals, we hypothesized that redox signaling in APCs mediates the activation of adipogenesis in response to a pro-inflammatory cytokine released by a neighbouring cell. The protein secretome collected from adipose explants facilitated differentiation of APCs with an amplified response when the secretome derived from explants of HFD-fed mice. Proteomics analysis revealed that the adipose secretome of HFD-fed animals was enriched in pathways involved in immune cell activation and identified IL-6 as central to differentially regulated proteins. Analysis of cytokine levels in the adipose secretome revealed higher levels of pro-inflammatory cytokines including IL-6 and TNFα after HFD-feeding, with IL-6 markedly more abundant than TNFα. Under normal conditions, exposure of APCs to IL-6, but not TNFα, enhanced adipogenesis. Neutralization of IL-6 had no impact on APC differentiation under normal conditions but blunted the potentiation of adipogenesis with exposure to the secretome of HFD-fed mice, the latter effect abolished when ROS were scavenged. Together, these findings suggest IL-6 to be a key adipokine involved in the differentiation of APCs under obesogenic conditions. These results are in agreement with an *in vivo* study by Asterholm *et al*, which demonstrated that inflammatory signaling in adipose tissue was required for adipogenesis and adaptive adipose remodeling in response to a HFD (23). In the study herein, we directly investigated the impact of TNFα on APC differentiation and found an inhibition of adipogenesis. This result is in agreement with several studies using 3T3-L1 cells and human APCs, which reported TNFα to decrease adipogenesis (24–26). Few studies have investigated the role of IL6 on adipogenesis which we found herein to increase adipogenesis *in vitro*.

Most studies in the obesity field focus on pathophysiology in the chronically obese state, while mechanisms driving adaptive adipose remodeling remain unclear. Emerging evidence suggests that similar to the recruitment of resident APCs to support normal adipocyte turnover, adipogenesis in the obese state may be activated to replace dead adipocytes that have reached their expansion limit (1). This may protect against adipose dysfunction that is characterized by excessive adipocyte hypertrophy, loss of insulin sensitivity in adipocytes, hypoxia, and fibrosis. Macrophages within the adipose tissue are adapted to the clearance of dead adipocytes and their lipid remnants *via* lysosomal biogenesis and lipid metabolism (6–9). Interestingly, a study using isotope labelling to measure adipocyte turnover found newly generated adipocytes in close proximity to crown-like structures, which are dead adipocytes surrounded by macrophages (13). Thus, APC differentiation may be induced by pro-inflammatory cytokines like IL-6 released from adipocytes undergoing cell death, recruited macrophages, or stimulated by free fatty acids released from dead adipocytes (27). This inflammatory event under homeostatic conditions is likely localized and short-lived, in contrast to the chronic inflammation that characterizes the adipose dysfunction accompanying severe obesity. Impaired adipogenesis has been implicated in obesity-induced metabolic disease and may result from APC dysfunction under conditions of chronic inflammation. Our recent work demonstrated that the capacity for adipogenesis is a key factor driving sex differences in the development of insulin resistance due to obesity (28). Future research could elucidate the role of IL-6-mediated crosstalk between APCs and neighbouring cells in the sex and depot-dependent regulation of adipogenesis.

---

**Supplementary Figure 1. Adipogenic Stimulus Increases the Number of Short Mitochondria.** (A) Mitochondrial length analyzed by category, short, intermediate, or elongated for APCs treated adipogenic stimulus or vehicle normalized to cell number per slide (B) Representative images of mitotracker stained APCs.

**Supplementary Figure 2. Conditioned Media Treatment Increases Adipogenesis.** (A) Eluted Oil Red O absorbance for cells treated with CD CM, HFD CM, or vehicle on D7 (n=4).

**Supplementary Figure 3. IL6 and TNF Identified as Upstream Regulators** (A) Network of IL6 as an upstream regulator of differentially expressed proteins found in our dataset in IPA. (B) Network of TNF as an upstream regulator of differentially expressed proteins found in our dataset in IPA.

## DECLARATIONS

### Ethics approval and consent to participate

All procedures involving animals were approved by the University of Calgary Animal Care Committee (Protocol #: AC21-0132) and conducted in accordance with guidelines by the Canadian Council on Animal Care Ethics.

### Availability of Data and Materials

Proteomics data will be made publicly available; all other data will be made available upon request.

### Author’s Contributions

JLW contributed to study design, collected and analyzed the data and wrote the manuscript. LGB contributed to data collection. TBS contributed to data collection and analysis. SZY contributed to data analysis. PC contributed to acquiring live cell imaging. AD contributed to proteomic data collection and analyses. JAT contributed to conception of the study and experimental design and contributed to the writing and editing of this manuscript.

## Acknowledgements

We acknowledge the Live Cell Imaging Laboratory at the University of Calgary for support with mitochondrial imaging and analysis.

