## Supplementary figures and images for "Redox Signaling Mediates Differentiation of Adipose Progenitors in Response to Inflammatory Cytokines in the Adipose Tissue Secretome"

### Supplementary Figure 1

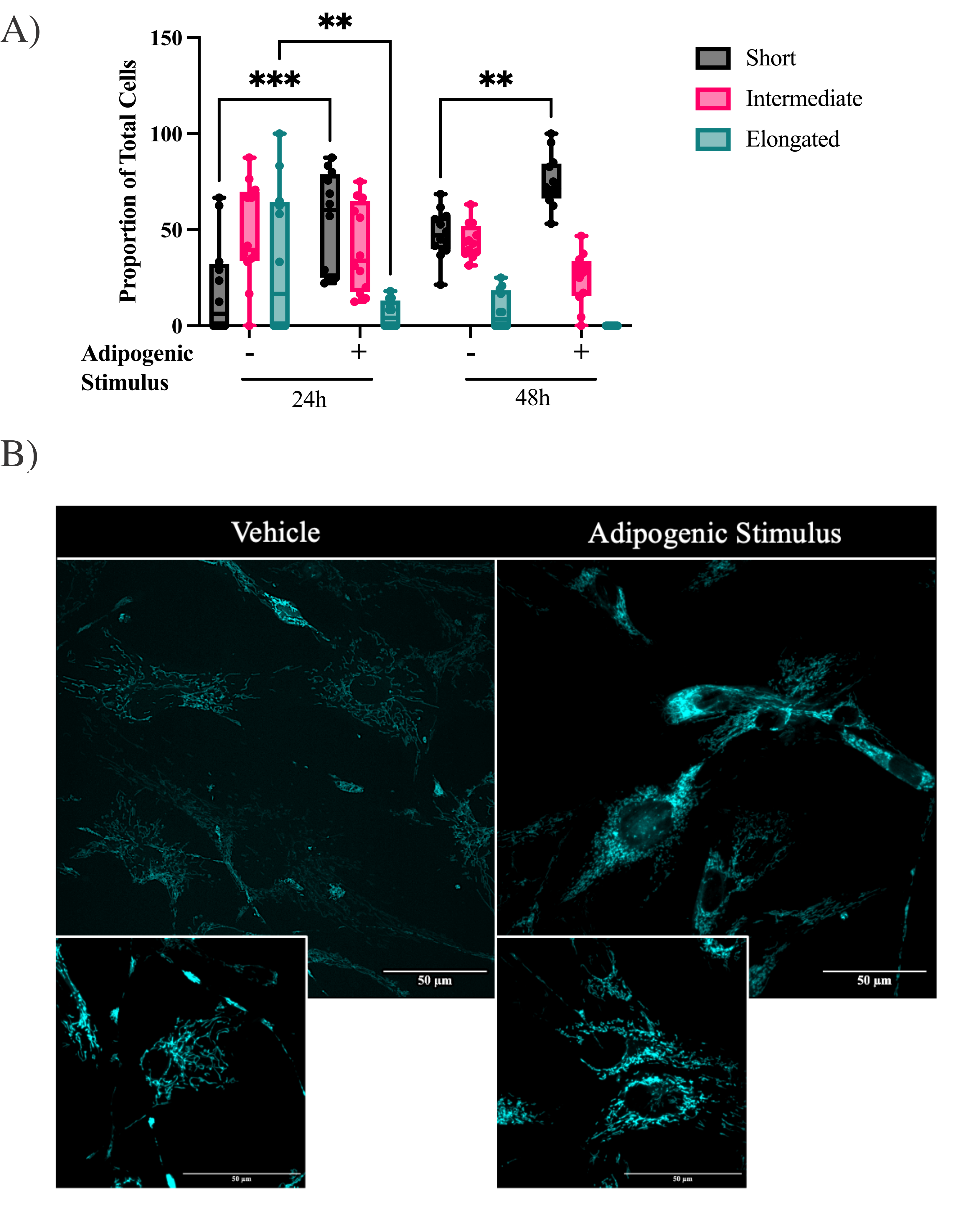

### Supplementary Figure 2

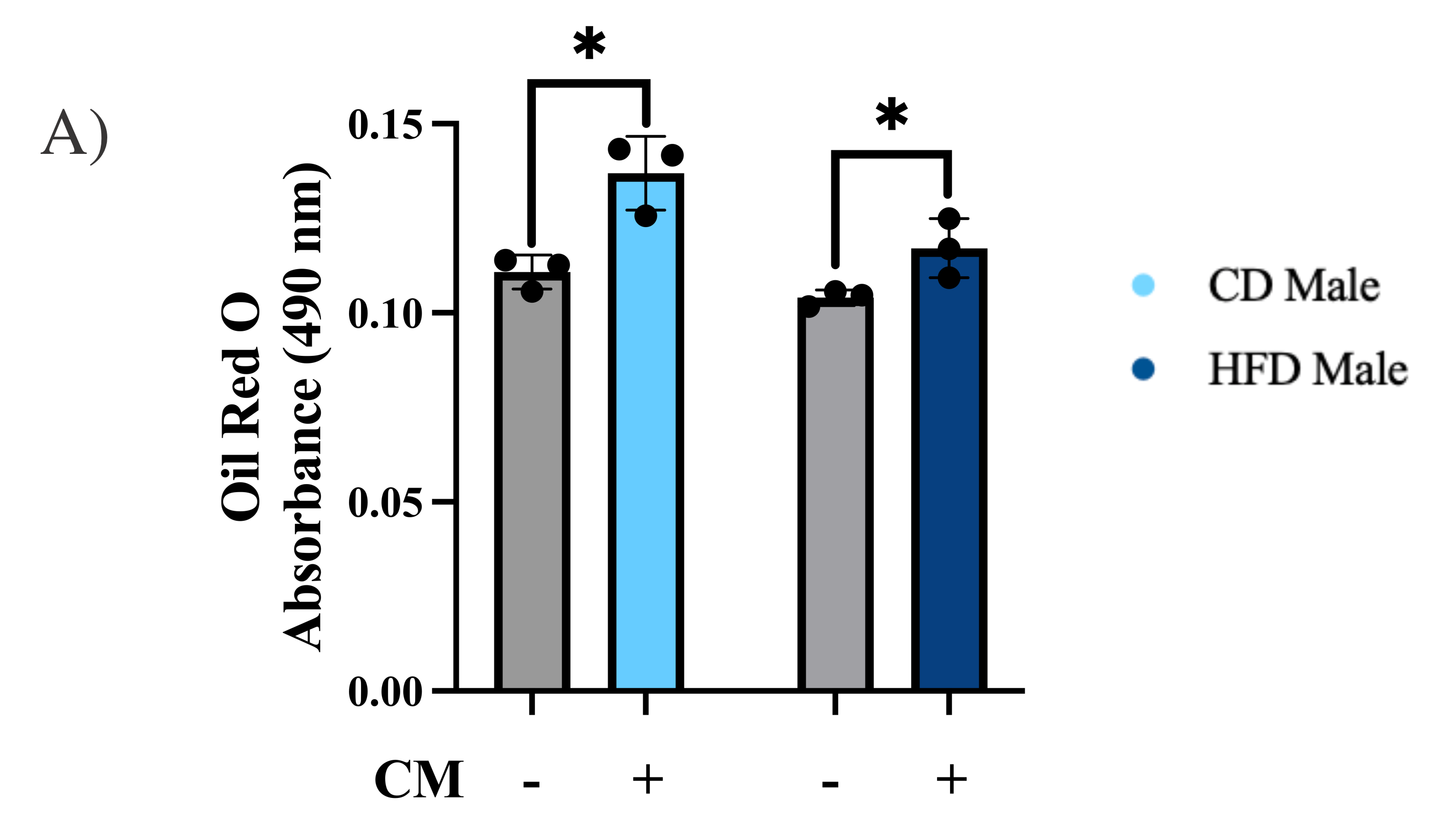

### Supplementary Figure 3

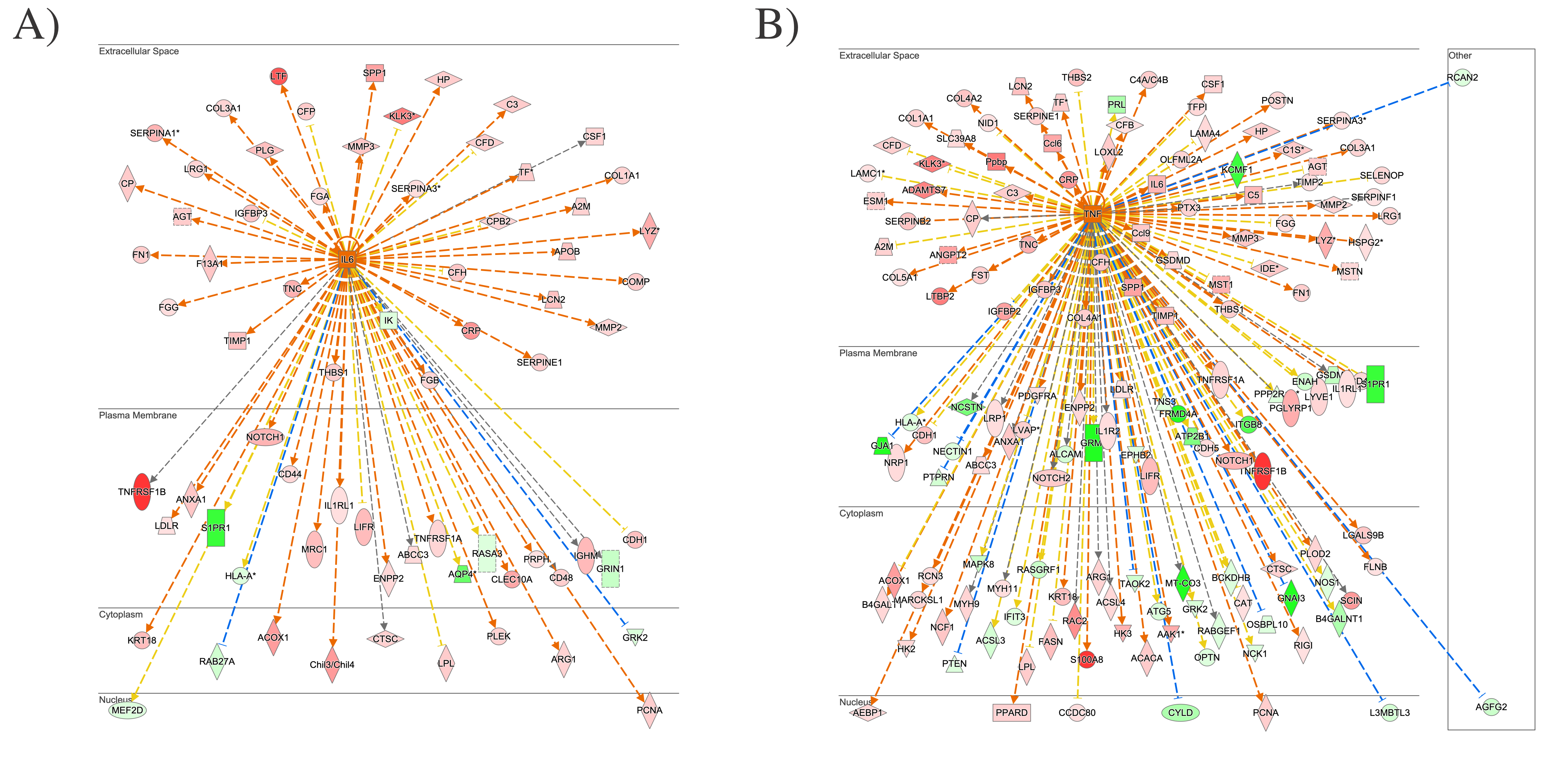
