## Supplementary Table 1 for "Redox Signaling Mediates Differentiation of Adipose Progenitors in Response to Inflammatory Cytokines in the Adipose Tissue Secretome"

### Slide 1
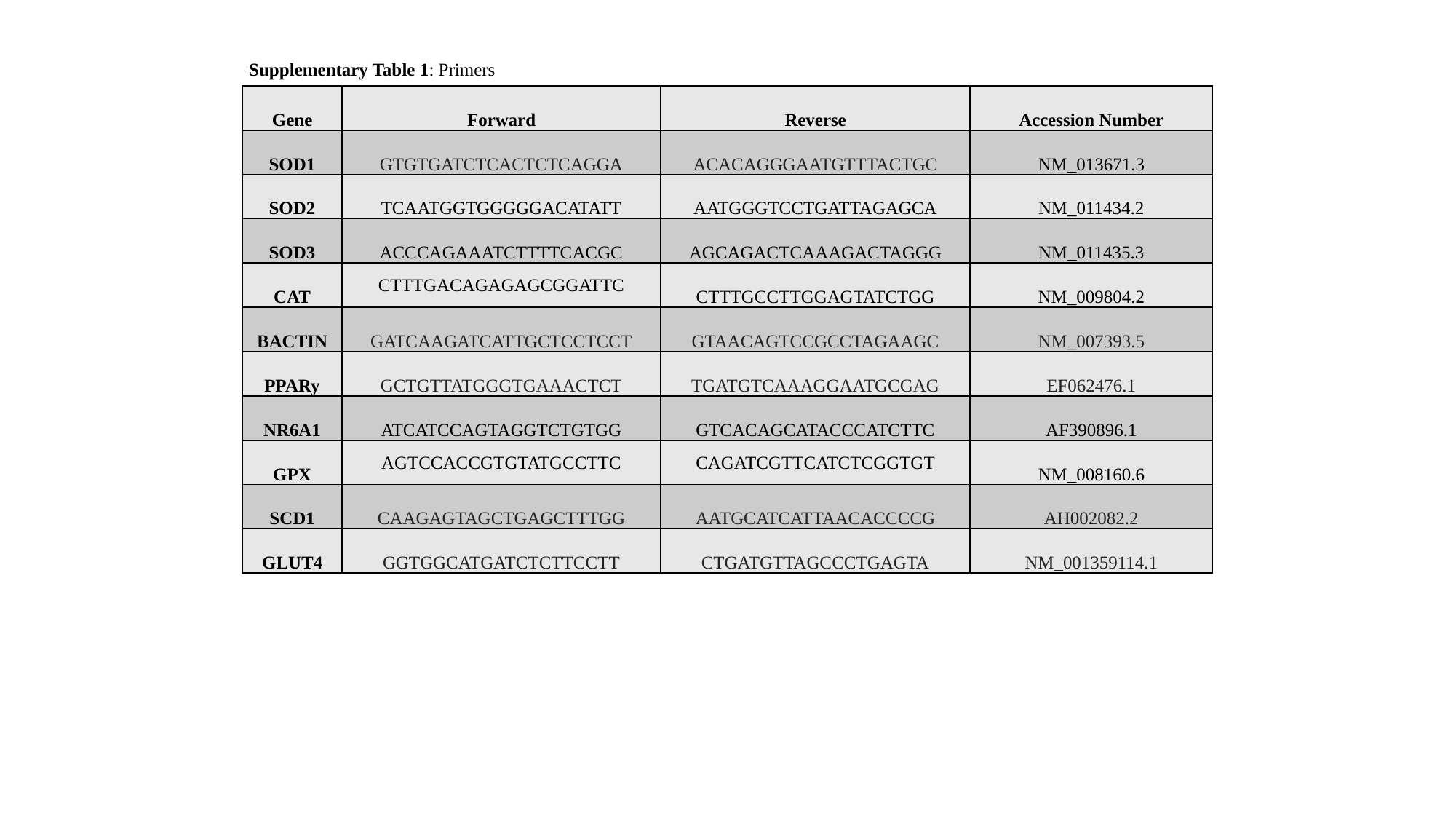

Supplementary Table 1: Primers
| Gene | Forward | Reverse | Accession Number |
| --- | --- | --- | --- |
| SOD1 | GTGTGATCTCACTCTCAGGA | ACACAGGGAATGTTTACTGC | NM\_013671.3 |
| SOD2 | TCAATGGTGGGGGACATATT | AATGGGTCCTGATTAGAGCA | NM\_011434.2 |
| SOD3 | ACCCAGAAATCTTTTCACGC | AGCAGACTCAAAGACTAGGG | NM\_011435.3 |
| CAT | CTTTGACAGAGAGCGGATTC | CTTTGCCTTGGAGTATCTGG | NM\_009804.2 |
| BACTIN | GATCAAGATCATTGCTCCTCCT | GTAACAGTCCGCCTAGAAGC | NM\_007393.5 |
| PPARy | GCTGTTATGGGTGAAACTCT | TGATGTCAAAGGAATGCGAG | EF062476.1 |
| NR6A1 | ATCATCCAGTAGGTCTGTGG | GTCACAGCATACCCATCTTC | AF390896.1 |
| GPX | AGTCCACCGTGTATGCCTTC | CAGATCGTTCATCTCGGTGT | NM\_008160.6 |
| SCD1 | CAAGAGTAGCTGAGCTTTGG | AATGCATCATTAACACCCCG | AH002082.2 |
| GLUT4 | GGTGGCATGATCTCTTCCTT | CTGATGTTAGCCCTGAGTA | NM\_001359114.1 |
